# BAY 11-7082 potentiates select β-lactams to inhibit growth of methicillin-resistant *Staphylococcus aureus*

**DOI:** 10.64898/2026.01.13.699313

**Authors:** Victoria E. Coles, Patricia Reed, Mariana G. Pinho, Lori L. Burrows

## Abstract

β-lactams are an important class of antibiotics that target penicillin-binding proteins (PBPs) essential to bacterial cell wall synthesis. They are used to treat serious bacterial infections, including those caused by the versatile pathogen *Staphylococcus aureus*. Some strains carry the resistance gene, *mecA,* encoding a β-lactam-insensitive PBP2A that allows for peptidoglycan synthesis in the presence of β-lactams like methicillin, limiting treatment options. We previously identified BAY 11-7082 as having antibacterial activity against methicillin-resistant *S. aureus* (MRSA) and showed that it re-sensitizes MRSA to β-lactams like penicillin G, making BAY 11-7082 and its analogues promising candidates for the development of an antibiotic adjuvant. Although the direct antibacterial mechanism of BAY 11-7082 remains undefined, here we aimed to better understand how it potentiates β-lactam activity. We tested BAY 11-7082 in combination with a variety of β-lactams and identified a subset that were synergistic in MRSA but not methicillin-susceptible *S. aureus*, suggesting they may directly or indirectly impact the function of PBP2A. Electron and fluorescence microscopy studies revealed that unlike other β-lactam adjuvants, such as those that target the biosynthesis of wall teichoic acids (WTAs), BAY 11-7082 failed to impact cell division, disrupt PBP2 localization, or activate a sensitive reporter of cell wall damage. However, the production of WTA was necessary for BAY 11-7082-β-lactam synergy. Together these data suggest its mechanism of β-lactam potentiation is distinct from known compounds, making BAY 11-7082 a useful tool to better understand the complexity of resistance in MRSA.

## INTRODUCTION

β-lactams are the most widely used class of antibiotics. They target penicillin-binding proteins (PBPs), essential enzymes that crosslink and tailor peptide stems to connect adjacent glycan strands during peptidoglycan synthesis. The broad spectrum and low toxicity of β-lactams contribute to their importance in the treatment of infections; however, their utility is decreased in the presence of β-lactamases, enzymes that bind and inactivate β-lactams. In such cases, β-lactams can be co-administered with a β-lactamase inhibitor to extend their usability ^1^. This is one of the few clinically validated examples of an antibiotic adjuvant, a therapeutic that enhances the effectiveness of an existing antibiotic ^2^. Some strains of the pathogen *Staphylococcus aureus* can carry the *mecA* gene that encodes for PBP2A, a β-lactam-insensitive PBP capable of crosslinking peptidoglycan even in the presence of a β-lactam, making these organisms especially difficult to treat ^3^. Despite the global dissemination of methicillin-resistant strains of *S. aureus* (MRSA), no antibiotic adjuvants that rescue the activity of β-lactams in *mecA*-expressing strains have yet been approved.

While PBP2A is required for β-lactam resistance in *S. aureus*, it is not sufficient; there are several auxiliary factors that contribute to resistance, representing possible targets for antibiotic adjuvants ^4,5^. These factors include genes involved in peptidoglycan biosynthesis (*murA-F*), assembly of the pentaglycine bridge that connects adjacent peptide stems (*femA,B,X*), cell division (*ftsA,Z*), and protein secretion (*spsB*)^5^. Inhibitors of FtsZ, an essential cell division protein that plays a role in PBP2 localization, and the type I signal peptidase SpsB, which is required for protein secretion through the Sec and TAT systems, have been identified ^6-9^. Other adjuvants target the biosynthesis of wall teichoic acids (WTAs), glycopolymer chains anchored to the peptidoglycan ^10^.

While not essential for *S. aureus* viability, WTAs stabilize the cell envelope and protect peptidoglycan from degrading agents like lysozymes ^11^. Importantly, WTAs have been implicated in proper PBP localization, which is required for PBP2A function. *S. aureus* strains lacking the WTA biosynthetic machinery have abnormal PBP4 localization, resulting in decreased peptidoglycan crosslinking ^12,13^. WTA biosynthesis in *S. aureus* and certain strains of *Bacillus subtilis* is carried out by proteins encoded by the *tar* genes (**Figure 1**) ^14^. The first enzyme in this pathway, TarO, reversibly catalyzes the transfer of GlcNAc phosphate to undecaprenyl phosphate. TarO inhibitors – including tunicamycin, ticlopidine, and tarocins – are nonlethal on their own but sensitize MRSA to certain β-lactams ^10,15,16^. The late stage WTA biosynthesis genes *tarBDI’KS* are auxiliary factors that when mutated, deleted, or depleted, restore the activity of β-lactams, but inhibitors of the flippase TarG, such as targocil, do not synergize with β-lactams and are lethal on their own ^17-20^. This phenotype is likely due to depletion of key precursors such as undecaprenyl phosphate that are also needed for peptidoglycan synthesis, as it can be alleviated by inhibiting the early-stage biosynthesis proteins TarO or TarA ^21,22^. By leveraging this conditionally lethal phenotype, Farha *et al.* and El-Halfawy *et al.* identified the β-lactam adjuvants clomiphene and MAC-545496 ^21,23^. Clomiphene, which has antibacterial activity on its own, antagonizes targocil by inhibiting undecaprenyl diphosphate synthase (UppS), limiting the availability of undecaprenyl phosphate ^21^. MAC-545496 inhibits the response regulator GraR, which regulates the *dltABCD* operon responsible for installing a *D*-alanine ester on WTA ^23,24^.

**Figure 1.**
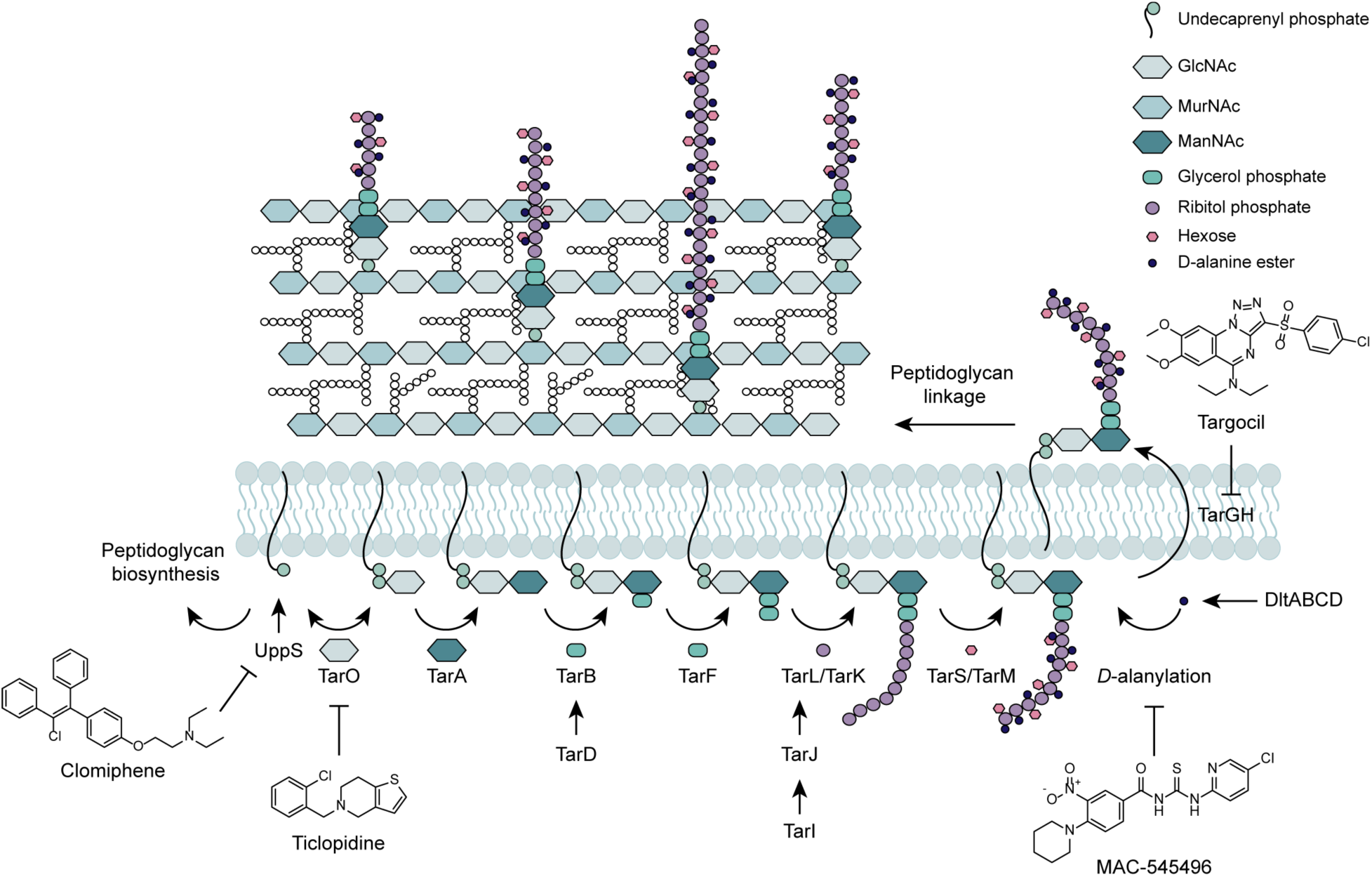
Wall teichoic acid biosynthetic pathway in *S. aureus*. Adapted from Swoboda *et al*. and Pasquina *et al*.^14,25^. Structures of chlomiphene, ticlopidine, MAC-545496, and targocil are shown as examples of UppS, TarO, *D*-alanylation, and TarG inhibitors, respectively.

We previously used a novel biofilm-based screening approach to identify compounds with sub-inhibitory antibiotic activity ^26^. We showed that the anti-inflammatory compound BAY 11-7082 had activity against a wide range of pathogens, with most potent activity against Gram-positive species including MRSA ^27^. We were unable to identify its specific antimicrobial mechanism of action, and concluded that its activity may be multifaceted. However, our discovery that it re-sensitizes MRSA to β-lactams like penicillin G motivated further investigation of this compound and its synthetic analogues as potential antibiotic adjuvants.

## RESULTS

### BAY 11-7082 synergizes with specific β-lactams

To probe the mechanism of action of BAY 11-7082, we previously examined the spectrum of antibiotics potentiated by the compound. We conducted checkerboard assays against the MRSA strain USA300 using a variety of classes, including aminoglycosides, tetracyclines, thiopeptides, glycopeptides, rifamycins, and β-lactams ^27^. Interestingly, while we observed synergy (as defined by a fractional inhibitory concentration index or FICI <0.5) ^28^ with the β-lactams penicillin G (FICI=0.19) and piperacillin (FICI=0.25), we saw no synergy with methicillin (FICI=0.56), to which *S. aureus* USA300 is intrinsically resistant. To further dissect this difference, we expanded the scope of β-lactams tested (**Figure 2, Figure S1**). We conducted checkerboard assays with MRSA strain USA300, which has *mecA* (**Figure 2B, Figure S2**), as well as methicillin-sensitive *S. aureus* (MSSA) strain ATCC 29213, which does not, to understand the potential role of PBP2a (**Figure 2C, Figure S3**).

**Figure 2.**
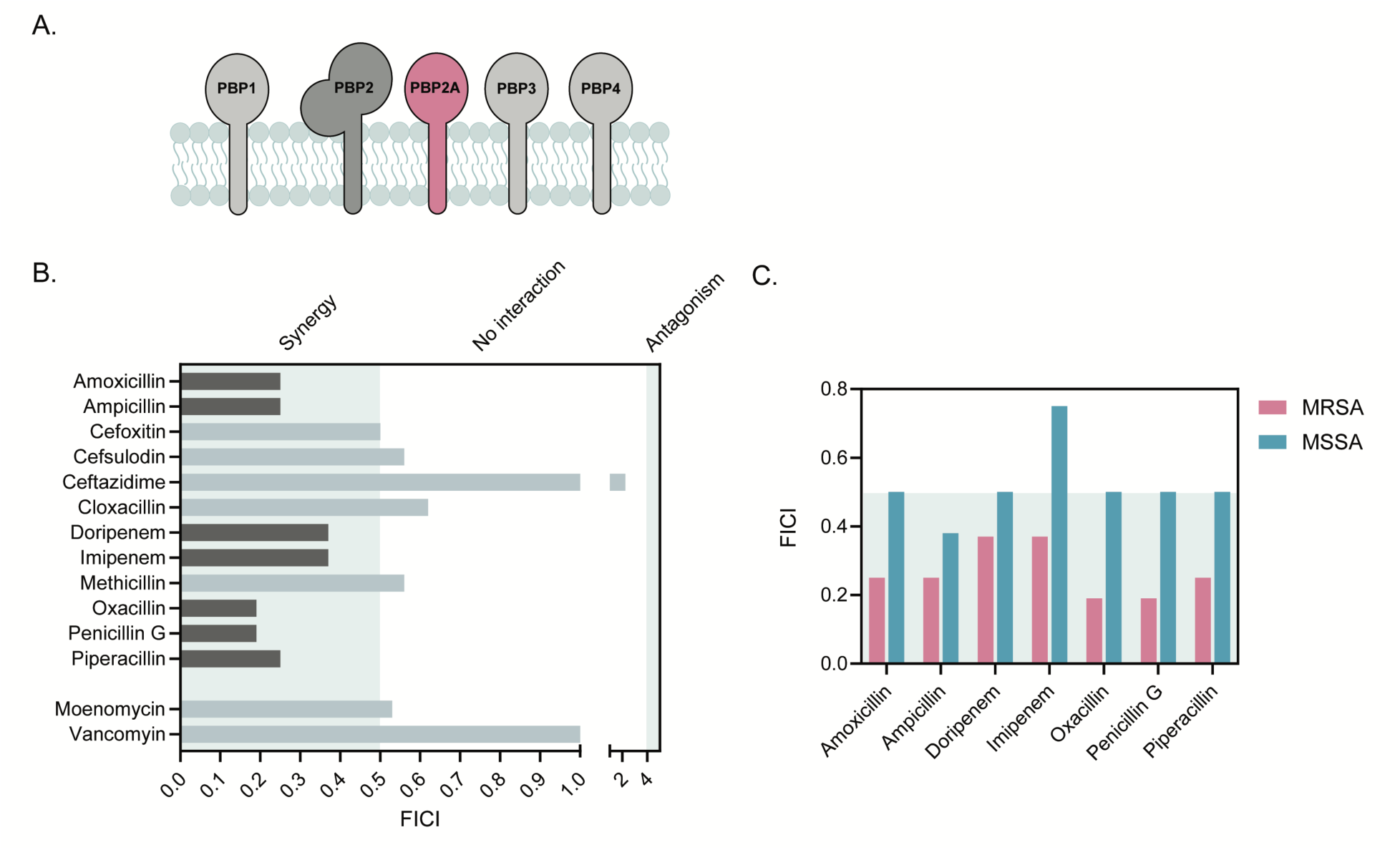
BAY 11-7082 synergizes with specific β-lactams. **A.** MRSA expresses five PBPs, PBP1, PBP2, PBP2A (pink), PBP3, and PBP4. **B.** FICI values calculated based on MRSA USA300 grown in the presence of BAY 11-7082 and another antibiotic. FICI values <0.5 represent synergy, values between 0.5-4 represent no interaction, and values >4 represent antagonism. **C.** FICI values for MRSA USA300 (pink) and MSSA ATCC 29213 (blue) grown in the presence of BAY 11-7082 and a β-lactam that synergizes with BAY 11-7082 in MRSA. Data represent an average of three independent experiments performed in triplicate.

Based on FICI values, BAY 11-7082 synergized with amoxicillin, ampicillin, doripenem, imipenem, oxacillin, piperacillin, and penicillin G, but not cloxacillin or methicillin (**Figure 2B**), and synergy was reduced or eliminated in the MSSA strain (**Figure 2C**). We also explored BAY 11-7082’s ability to potentiate two non-β-lactam antibiotics that target cell wall biosynthesis (**Figure 2B**). BAY 11-7082 failed to synergize with vancomycin, which prevents transglycosylation and transpeptidation by binding the terminal amino acids of lipid II, or moenomycin, a glycosyltransferase inhibitor that also synergizes with β-lactams.

The loss of interaction for most of the synergistic beta lactam combinations in MSSA suggested that BAY11-7082 may directly or indirectly impact the function of MRSA-specific PBP2A. PBP2 is the only bifunctional PBP in *S. aureus*, with both transpeptidase and glycosyltransferase activity and, along with PBP1, is essential. PBP1 and PBP3 are transpeptidases that coordinate with the SEDS (shape, elongation, division, and sporulation) proteins FtsW and RodA to facilitate peptidoglycan synthesis during cell division and elongation, respectively ^29^, while PBP2, PBP4, and PBP2A function cooperatively to produce highly crosslinked peptidoglycan strands ^30,31^. The transpeptidase domain of PBP2A can crosslink peptidoglycan in the presence of β-lactams; however, it lacks a glycosyltransferase domain, necessitating coordination with the glycosyltransferase domain of PBP2 ^31,32^. We hypothesized that BAY 11-7082 might inhibit the glycosyltransferase activity of PBP2, similar to the antibiotic moenomycin, or result in PBP2 mislocalization, interrupting coordination with PBP2A and re-sensitizing MRSA to β-lactams. To determine if BAY 11-7082 acts similarly to moenomycin, we used an MRSA 15981 mutant previously selected for increased resistance to BAY 11-7082 (B13), along with a mutant selected for increased resistance to a structural analogue 3-(phenylsulfonyl)-2-pyrazinecarbonitrile (PSPC) (P11) ^27^. While the minimum inhibitory concentration (MIC) of moenomycin was two-fold higher in both mutants compared to the parent strain, neither mutant displayed cross-resistance, suggesting that BAY 11-7082 and moenomycin act via distinct mechanisms (**Figure S4**).

### Wall teichoic acid synthesis may play a role in BAY 11-7082–β-lactam synergy

Compounds that inhibit FtsZ and WTA biosynthesis are proposed to synergize with certain β-lactams against MRSA by causing PBP mislocalization ^6,8,10,15,16^, leading us to ask if BAY 11-7082 acts on one of these pathways. To explore BAY 11-7082’s impact on WTA biosynthesis, we measured activity against a Δ*tarO* mutant of *S. aureus* USA300 which lacks the enzyme that catalyzes the first biosynthetic step ^10^(**Figure 1**). As expected, the inhibitory concentration of penicillin G dropped eight-fold in the Δ*tarO* strain (**Figure 3A**). The inhibitory concentration of BAY 11-7082 for the Δ*tarO* mutant was similar to that for the parent strain, suggesting that BAY 11-7082’s direct antibacterial activity is not dependent on WTA biosynthesis (**Figure 3A**).

**Figure 3.**
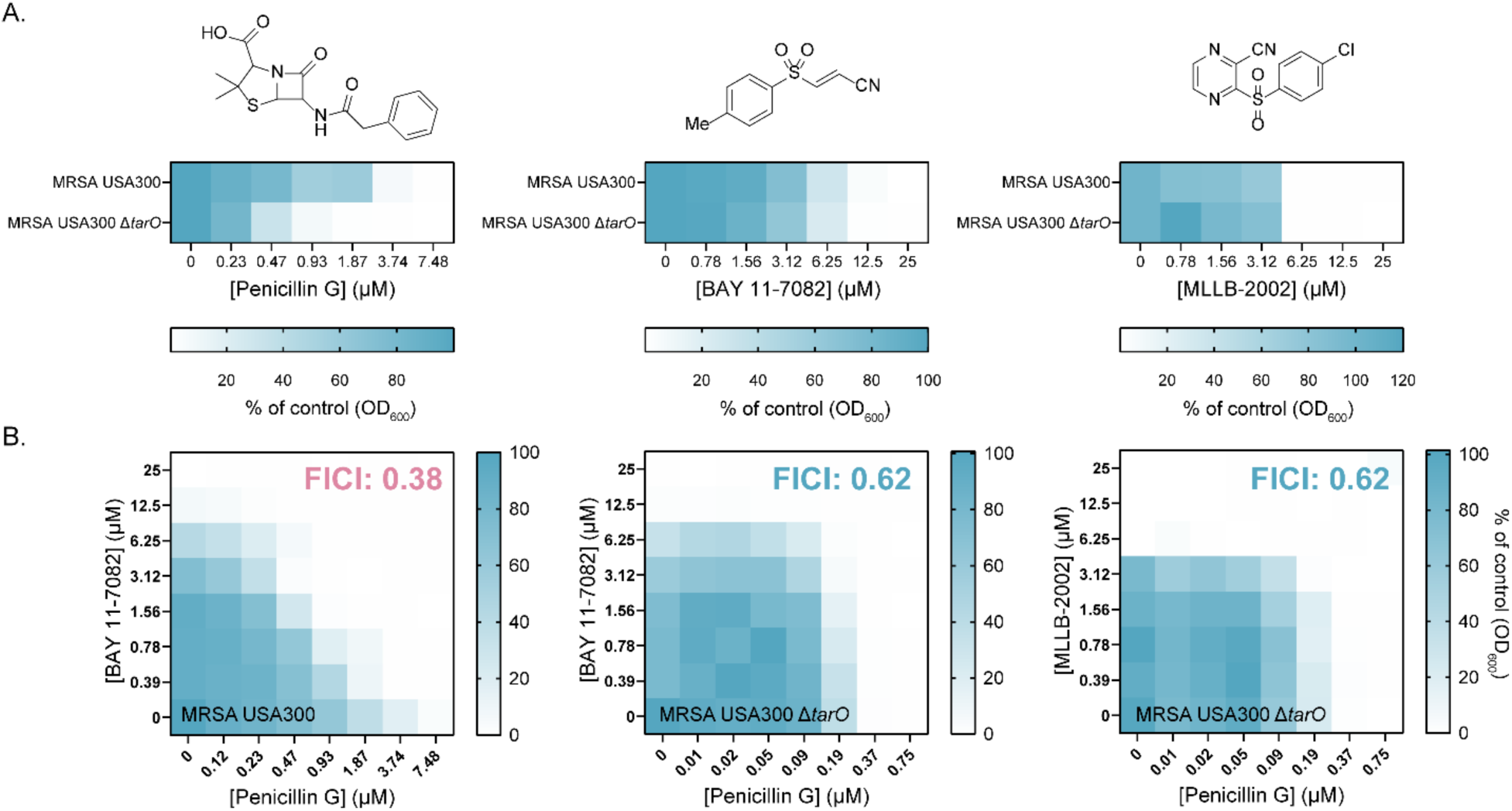
WTA synthesis is required for BAY 11-7082 to potentiate penicillin G activity. **A.** Planktonic growth (optical density at 600 nm as a percent of the vehicle control) of *S. aureus* USA300 and a Δ*tarO* mutant grown in the presence of increasing concentrations of penicillin G (left), BAY 11-7082 (middle), or the analogue MLLB-2002 (right) (µM). **B.** Checkerboards of *S. aureus* USA300 (left) or a Δ*tarO* mutant (middle and right) grown with increasing concentration of penicillin G and BAY 11-7082 or MLLB-2002, where synergy (FICI < 0.5) is represented in pink text while indifference (FICI ≥ 0.5) is in blue. Data represent an average of three independent experiments.

However, BAY 11-7082 no longer potentiated penicillin G activity in the Δ*tarO* strain (FICI=0.62) (**Figure 3B**), suggesting that while BAY 11-7082’s antibacterial activity is unrelated to WTA biosynthesis, the pathway impacts its ability to synergize with β-lactams. To confirm that these phenotypes remained consistent for structural analogues of BAY 11-7082, we also measured the activity of the potent chlorinated PSPC-analogue MLLB-2002 (**Figure 3**), with similar results.

These phenotypes are similar to those reported for cells treated with the UppS inhibitor clomiphene (**Figure 1**), which also sensitizes MRSA to β-lactams but retains its activity in strains lacking WTA ^21^. This observation raised the question of whether BAY 11-7082 inhibits an upstream enzyme common to both WTA and peptidoglycan biosynthesis. Inhibitors that deplete shared substrates of WTA and peptidoglycan synthesis such as undecaprenyl phosphate can antagonize activity of the TarG inhibitor targocil by limiting resources available for WTA biosynthesis, reducing its effect on the cell ^21^. Unexpectedly, BAY 11-7082 did not impact targocil activity (FICI=1.5) (**Figure 4A**), suggesting BAY 11-7082 does not decrease WTA biosynthesis. We also measured the impact of BAY 11-7082 treatment on susceptibility to lysostaphin, an endopeptidase capable of cleaving the pentaglycine bridge in *S. aureus* peptidoglycan, since strains lacking WTA are more sensitive than wild type (**Figure 4B, D**). Lysostaphin sensitivity was not altered in the presence of BAY 11-7082 (FICI=0.53) (**Figure 4C**). Taken together, these results suggest BAY 11-7082 does not impact WTA levels.

**Figure 4.**
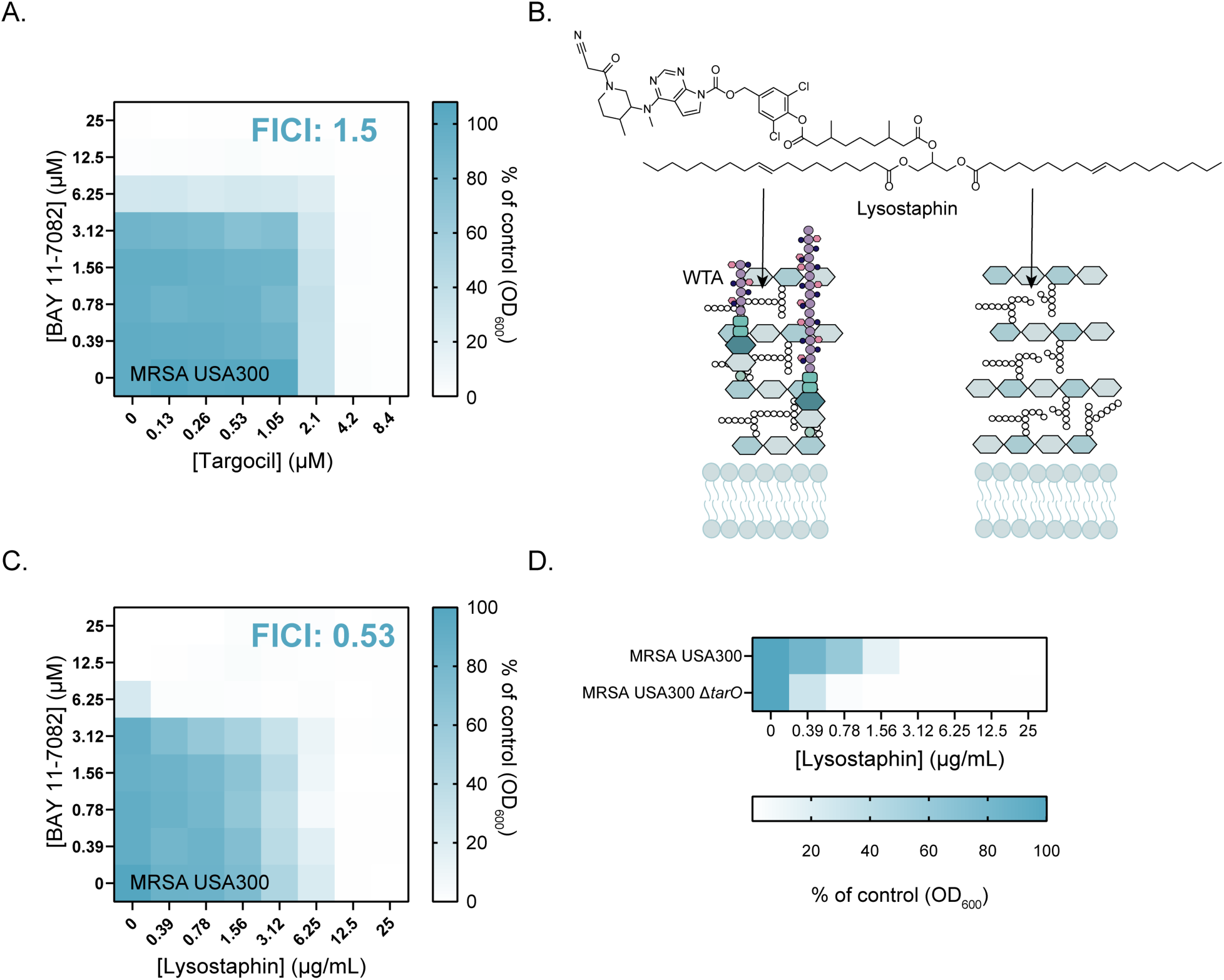
BAY 11-7082 does not impact WTA levels. **A.** Checkerboard of *S. aureus* USA300 grown with increasing concentration of BAY 11-7082 and the TarG inhibitor targocil (µM). **B.** Lysostaphin is an endopeptidase that cleaves the pentaglycine bridge in *S. aureus* peptidoglycan. Cells producing WTA are less sensitive (left) compared to those lacking WTA (right). **C.** Checkerboard of *S. aureus* USA300 grown with increasing concentration of BAY 11-7082 (µM) and lysostaphin (µg/mL). **D.** Planktonic growth (optical density at 600 nm as a percent of the vehicle control) of *S. aureus* USA300 and a Δ*tarO* mutant grown in the presence of increasing concentrations of lysostaphin (µg/mL). Data is based on three independent experiments preformed in triplicate. FICI values ≥ 0.5 represent indifference, and checkerboards are based on an average of three independent experiments.

### WTA glycosylation is not required for BAY 11-7082-β-lactam synergy

While inconsistent with inhibition of TarO, lack of synergy with lysostaphin is consistent with inhibition of TarS, the glycosyltransferase responsible for installing a β-*O*-GlcNAc on *S. aureus* WTA ^17^. Mutants lacking *tarS* are viable but hypersensitive to a wide range of β-lactams ^17^. To explore whether WTA glycosylation impacted BAY 11-7082’s ability to synergize with β-lactams, we made use of the Nebraska transposon library, a set of MRSA USA300 mutants containing single transposon insertions in non-essential genes ^33^. A mutant with a transposon insertion in *tarS* had similar susceptibility to BAY 11-7082 as the parent strain (**Figure 5A**), and remained susceptible to a combination of BAY 11-7082 and penicillin G (FICI = 0.37) (**Figure 5B**), indicating that while WTAs are required for synergy, their glycosylation is not.

**Figure 5.**
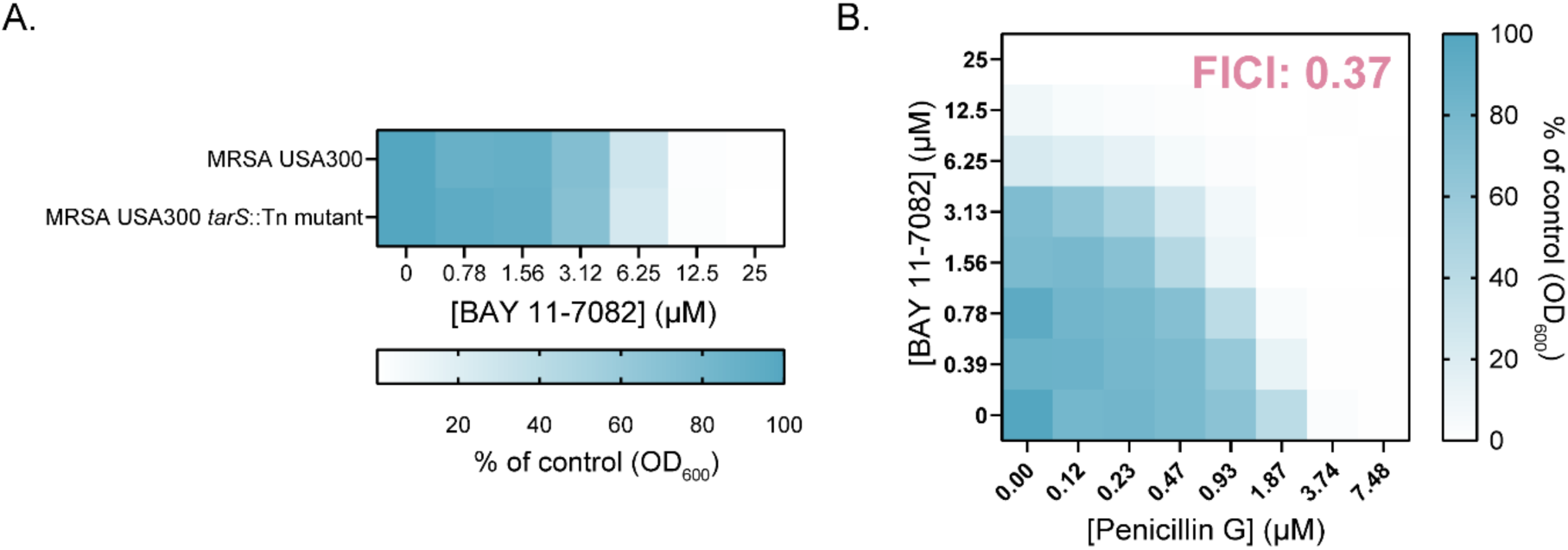
WTA glycosyltransferase TarS is not required for BAY 11-7082-PenG synergy. **A.** Planktonic growth (optical density at 600 nm as a percent of the vehicle control) of *S. aureus* USA300 and a *tarS* transposon mutant grown in the presence of increasing concentrations of BAY 11-7082 (µM). **B.** Checkerboard of a *tarS* transposon mutant of *S. aureus* USA300 grown with increasing concentration of penicillin G and BAY 11-7082 (µM) where synergy (FICI < 0.5) is indicated in pink text. Data represent an average of three independent experiments.

### BAY 11-7082 does not impact cell division

To further probe the mechanism of BAY 11-7082-β-lactam synergy and how WTAs may be involved, we examined whether BAY 11-7082 treatment impacts cell division. Septal abnormalities and division defects occur in cells treated with inhibitors of auxiliary factors like FtsZ and are also characteristic of cells lacking WTA, lipoteichoic acids (LTAs), or *D*-alanine esters ^15,24^. Cells without functional TarS, however, have no morphological or division defects ^17^. We used transmission electron microscopy (TEM) to examine thin-sectioned untreated MRSA cells (**Figure 6A, Figure S5A**) and cells exposed to 8x or 16x MIC of BAY 11-7082 to look for division defects (**Figure 6D, Figure S5D-E**). As controls, we also imaged cells treated with targocil plus untreated Δ*tarO* cells (**Figure 6B-C, Figure S5B-C**). As expected, Δ*tarO* cells had a thinner outer layer and less well-defined edges compared to wild type, consistent with published phenotypes for cells lacking WTA, and targocil-treated cells were clustered together, indicating improper separation following division ^15,18^. In contrast, cells treated with BAY 11-7082 more closely resembled untreated wild-type cells with sharp, thick peripheries, suggesting an intact peptidoglycan layer rich in WTAs, septa formation along the midcell, and separation following division. These data suggest that BAY 11-7082 synergizes with β-lactams via a mechanism unlike those described previously for other WTA-targeting compounds.

**Figure 6.**
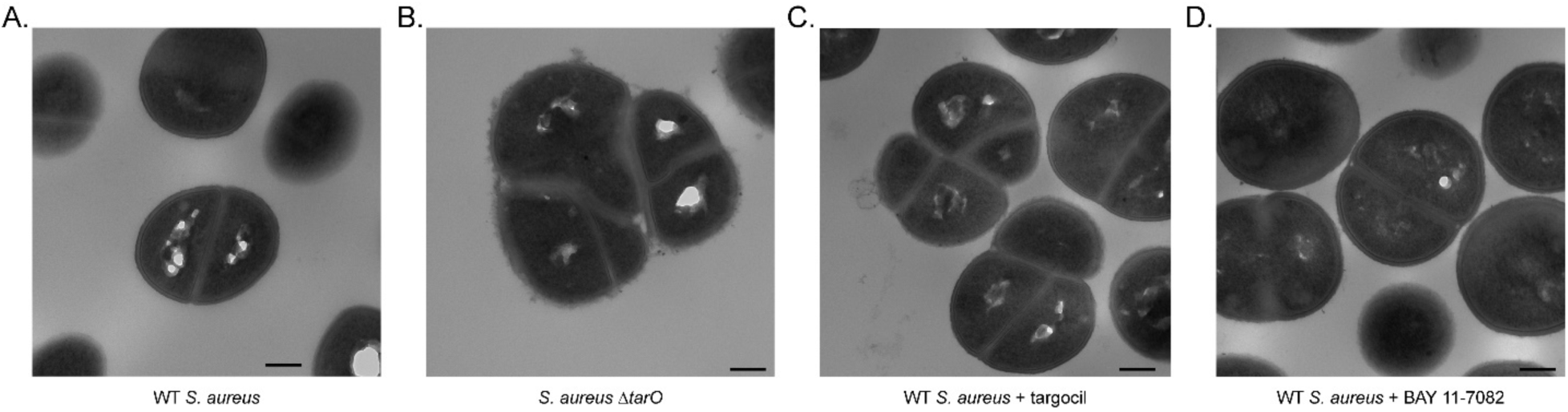
BAY 11-7082 treatment causes no morphological changes or septal abnormalities. **A.** Electron micrograph of wildtype (WT) untreated *S. aureus* USA300. **B.** *S. aureus* USA300 *ΔtarO* cells exhibit morphological defects including irregular septum formation. **C.** Wild type *S. aureus* USA300 treated with 33.6 µM targocil fail to separate following division. **D.** Wild type *S. aureus* USA300 treated with 50 µM BAY 11-7082 display no obvious morphological abnormalities. Scale bar = 200 nm. Additional images are available in Figure S5.

### BAY 11-7082 is unlikely to directly target peptidoglycan synthesis

Peptidoglycan synthesis inhibitors usually cause delocalization of PBP2 from the division septum, as this protein requires the presence of its substrate at the septum to localize correctly^34^. To look for potential mislocalization of PBP2 following BAY 11-7082 treatment, we incubated *S. aureus* strain COLsfGFP-PBP2, expressing a fluorescent derivative of PBP2, with and without BAY 11-7082. No PBP2 delocalization was observed (**Figure 7**). To further test if BAY 11-7082 was likely to target peptidoglycan synthesis, we used it to treat strain JE2 Pvra-mNG, where mNeonGreen expression is driven by the *vraTSR* promoter. The VraTSR system responds to cell wall damage and its expression is induced in the presence of antibiotics that target different steps of peptidoglycan synthesis, such as vancomycin, oxacillin, bacitracin, D-cycloserine or fosfomycin ^34-36^. Unlike the control compound vancomycin, BAY 11-7082 did not activate *vraTSR* promoter (**Figure 8**) suggesting that it does not target peptidoglycan synthesis.

**Figure 7:**
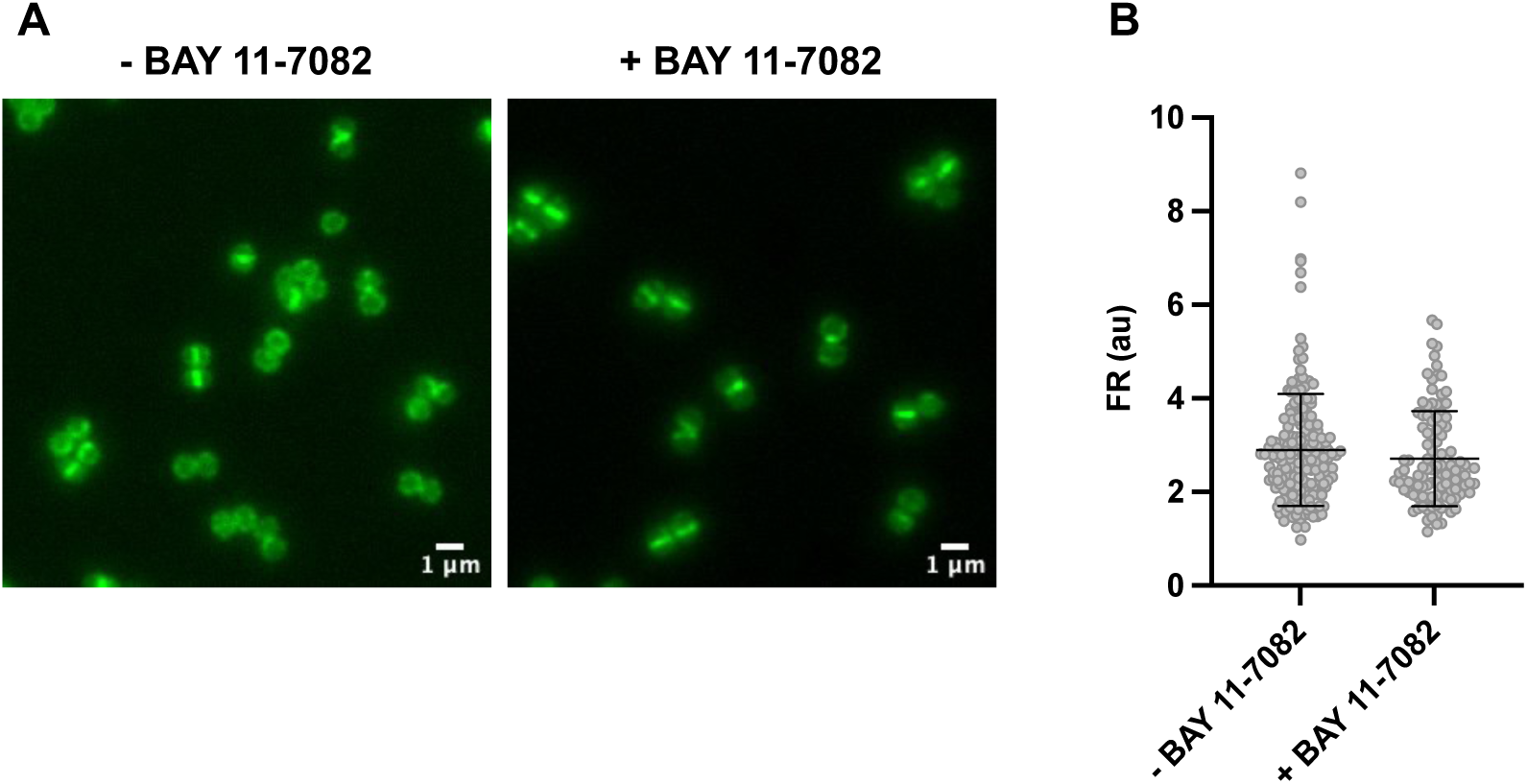
BAY 11-7082 treatment does not delocalize PBP2. **A.** Epifluorescence microscopy images of strain COL sfGFP-PBP2, expressing a fluorescent derivative of PBP2. sfGFP-PBP2 remained enriched at the septum in the presence of BAY 11-7082 (5X MIC). Scale bar = 1 μm. **B.** Quantification of the fluorescence ratio (FR) of sfGFP-PBP2 fluorescence at the septum versus the cell periphery in cells imaged as in panel **A.** Delocalization of PBP2 would result in lower FR, which was not observed in the presence of BAY 11-7082.

**Figure 8:**
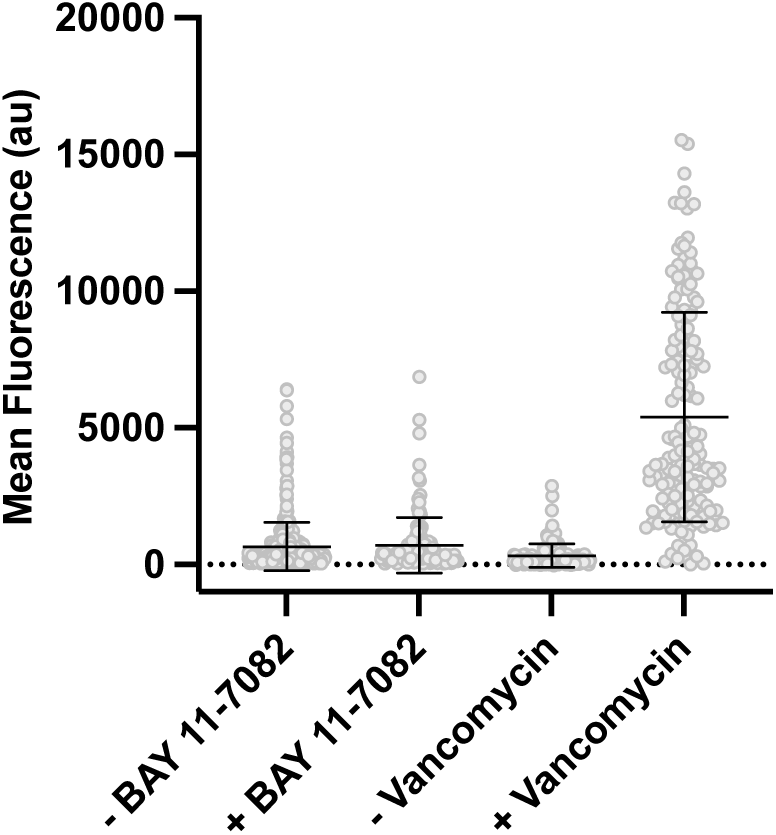
BAY 11-7082 failed to activate the VraTSR two-component system. Strain JE2 Pvra-mNG, which expresses mNeonGreen under the control of the *vraTSR* promoter, was incubated in the absence or presence of BAY 11-7082 (5X MIC) or vancomycin (10X MIC) and the mean fluorescence of cells was measured. The *vraTSR* promoter was activated in the presence of vancomycin, but not in the presence of BAY 11-7082.

## DISCUSSION

The anti-inflammatory compound BAY 11-7082 was first identified in a previous screen due to its broad-spectrum antibacterial activity ^26,27^, and further investigated here for its potential as a β-lactam adjuvant for MRSA. We first identified a subset of β-lactams potentiated by BAY 11-7082 (**Figure 2B**), and showed that synergy for most compounds was lost in an MSSA background, implying a potential impact on the function of PBP2A. These findings led us to speculate that BAY 11-7082 might interrupt the crucial coordination between the β-lactam insensitive transpeptidase domain of PBP2A and the glycosyltransferase domain of PBP2, which is required for proper cell wall synthesis in the presence of a β-lactam ^31^. This idea is consistent with affinity data showing that the synergistic subset can inhibit PBP2 ^6^. Interestingly, cloxacillin, which differs from oxacillin by the addition of a single chlorine atom (**Figure S1**), did not synergize with BAY 11-7082 (FICI = 0.62), while oxacillin did (FICI = 0.19) (**Figure 2B, Figure S2**), likely because cloxacillin, unlike oxacillin, is highly specific for PBP1 ^37^. PBP4 is a nonessential PBP with transpeptidase activity that, when inactivated, sensitizes cells to β-lactams specific for PBP2.^28^ Since PBP2 and PBP4 function cooperatively to crosslink glycan strands,^28^ we expected BAY 11-7082 to synergize with cefoxitin, a β-lactam with high affinity for PBP4. While combining BAY 11-7082 with cefoxitin resulted in a low FICI of 0.5 (**Figure 2B, Figure S2**), this interaction just exceeded the mathematical cut-off (FICI<0.5) for synergy.

Although we hypothesized that BAY 11-7082 may interrupt coordination between PBP2 and PBP2A, potentially through PBP2 mislocalization, this hypothesis was not supported by our fluorescence microscopy studies (**Figure 7**). Fluorescently-tagged PBP2 was enriched at the septum even in the presence of 5XMIC of BAY 11-7082. The lack of BAY 11-7082’s impact on PG synthesis was also supported by its inability to activate the VraTSR two-component system, unlike the control compound, vancomycin.

While BAY 11-7082 failed to synergize with the glycopeptide antibiotic vancomycin (**Figure 2B**), it did cause a slight reduction in sensitivity (**Figure S2**). This phenotype resembles the effect of a point mutation in the phosphorylation site of *walR*, which encodes the response regulator of WalKR, an essential two-component system that regulates autolysins involved in cleaving peptidoglycan during cell growth.^32^ Despite this similarity, *walR* mutants are not more susceptible than wild type to PBP2-targeting β-lactams like oxacillin,^32^ suggesting BAY 11-7082 acts via a different mechanism.

If BAY 11-7082 does not disrupt coordination of PBP2 and PBP2A, it could synergize with PBP2-targeting β-lactams by directly inhibiting the glycosyltransferase activity of PBP2, similar to the antibiotic moenomycin. We showed that BAY 11-7082 neither potentiated nor antagonized moenomycin activity (**Figure 2B, Figures S2-S3**), and cells with increased resistance to BAY 11-7082 or its structural analogue PSPC did not exhibit significantly increased resistance to moenomycin (**Figure S4**). Moenomycin resistance is rare, and few groups have identified moenomycin-resistant strains. Notably, in 2013 the Walker lab identified two single point mutations in the active site of the glycosyltransferase domain of PBP2 that each confer resistance to moenomycin, but they are not present in strains resistant to BAY 11-7082 ^27,38^, suggesting that if BAY 11-7082 interrupted the glycosyltransferase activity of PBP2, it would do so via an alternate mechanism.

Since several auxiliary factors are involved in proper PBP coordination, we wondered whether BAY 11-7082 acted on any of those. Many auxiliary factors are directly involved in peptidoglycan biosynthesis, but some have roles in other cellular processes, including the biosynthesis of WTAs ^10^. We showed BAY 11-7082 does not target WTA biosynthesis, since it and its synthetic analogues retain potency in a Δ*tarO* mutant (**Figure 3A**). We did, however, find that WTA biosynthesis was required for BAY 11-7082-β-lactam synergy (**Figure 3B**). We also considered that BAY 11-7082’s direct antibacterial target is distinct from the target that potentiates β-lactam activity. This idea is consistent with our previous work that suggested BAY 11-7082 could have multiple cellular targets, and with its activity against Gram-negative species including *Pseudomonas aeruginosa*, *Escherichia coli*, and *Klebsiella pneumoniae* ^27^.

Many compounds that synergize with β-lactams against MRSA have been identified in the literature, but no adjuvants have made it to the clinic. For example, the β-lactam cefoxitin itself potentiates the activity of other β-lactams by inhibiting PBP4, which acts cooperatively with PBP2 and PBP2A ^30,39^. The lipopeptide daptomycin and natural flavonoid epicatechin gallate, which both target the cytoplasmic membrane, synergize with oxacillin ^40,41^. Epicatechin gallate causes PBP2 mislocalization by intercalating in the cytoplasmic membrane and reducing *D*-alanylation of WTAs; however, unlike BAY 11-7082, it induces morphological changes, interrupts proper cell separation following division, and decreases lysostaphin susceptibility ^40,42^. MAC-545496 synergizes with β-lactams, including oxacillin, by inhibiting the response regulator GraR, which regulates *dltABCD*, among other genes, but antagonizes the TarG inhibitor targocil, similar to the UppS inhibitor clomiphene ^21,23^. FtsZ inhibitors cause PBP2 mislocalization, but also induce division defects and cell wall invaginations ^6,8^.

Due to differences in phenotype caused by previously identified adjuvants, we expect that BAY 11-7082 acts in a unique way, possibly by targeting auxiliary factors that play a role in β-lactam resistance that have not previously been targeted. LTAs, unlike WTAs, are linked to the cell membrane, synthesized via a separate pathway, and are essential ^43^. A transposon insertion in the LTA synthase gene *ltaS* causes increased sensitivity to β-lactams, suggesting those polymers also play a role in modifying β-lactam resistance ^44^. While inhibitors of LTA biosynthesis have not been identified, an *ltaS* transposon mutant is resistant to inhibitors of the signal peptidase SpsB, which regulates LtaS ^44,45^. However, strains lacking LTA were reported to be enlarged with improper septation and separation, unlike cells treated with BAY 11-7082 (**Figure 6D**)^24^. Cells lacking the auxiliary factor TarS are morphologically similar to those treated with BAY 11-7082 ^17^, but synergy was maintained in a *tarS* transposon mutant, indicating it is unlikely to be a target (**Figure 5B**). Since BAY 11-7082 is unlikely to inhibit these previously characterized auxiliary factors, it may impact the function of a determinant of β-lactam resistance that has not yet been characterized.

In conclusion, BAY 11-7082 and its structural analogues synergize with select β-lactams against MRSA, restoring the activity against this challenging pathogen of inexpensive and readily available compounds such as penicillin G. While WTA biosynthesis is required for this synergistic interaction, our data suggest that BAY 11-7082 does not target WTA nor the synthesis of shared precursors like undecaprenyl phosphate. Unlike previously characterized adjuvants, BAY 11-7082 does not cause aberrant cell division or separation. Together, the data suggest that BAY 11-7082’s mechanism of potentiation makes it unique among known β-lactam adjuvants, and thus it will be useful as a tool to better understand PBP2A-mediated β-lactam resistance.

## METHODS

### Bacterial strains, culture conditions, and chemicals

For synergy studies we used *S. aureus* USA300 (MRSA) and *S. aureus* ATCC 29213 (MSSA). *S. aureus* USA300 and the transposon mutant presented in Figure 5 were from the Nebraska transposon mutant library ^33^. *S. aureus* USA300 *ΔtarO* was kindly gifted by Eric Brown’s laboratory ^10^. BAY 11-7082 and PSPC resistant mutants of *S. aureus* 15981 (MRSA) were generated as previously reported ^27^.

Plasmid pBCB60 was constructed by digesting low-copy-number shuttle vector pCN34 ^46^ with BamHI/XbaI and inserting the gene encoding mNeonGreen, codon-optimized for *S. aureus* ^47^.To construct the *vraTSR* promoter fusion, a 704-bp DNA fragment containing the *vraTSR* promoter region was amplified from JE2 ^33^ genomic DNA using primers 8301 (GCGCGCGGATCCCGGTGCTATTTCTGCGCC) and 8302 (GCGCGCGGATCCTTATAATAAGTTTTAAAATACCAAATGCGC), digested with BamHI, and cloned into BamHI digested pBCB60, upstream of the mNeonGreen gene, resulting in plasmid pPvra-mNG, confirmed by DNA sequencing. Plasmid pPvra-mNG was electroporated into the *S. aureus* RN4220 strain; the resulting strain was named RN Pvra-mNG. Strain JE2 Pvra-mNG was constructed by transducing pPvra-mNG from RN Pvra-mNG into JE2 using phage 80α, as previously described ^48^. Strain COL sfGFP-PBP2 was constructed by introducing the plasmid pBCBPM061^8^ into RN4220 by electroporation, resulting in strain RN pBCBPM061, then transducing the plasmid to strain COL ^49^ using phage 80α ^48^. Integration and excision of the plasmid into the chromosome was performed as previously described ^50^.

Bacterial cultures were grown in Mueller-Hinton broth (MHB). Solid media was solidified with 1.5% agar. BAY 11-7082 and its structural analogue MLLB-2002 were synthesized as previously reported and stored as 100 µM stocks in DMSO at -20 °C ^27^. Stock solutions of amoxicillin (Sigma, prepared in DMSO), ampicillin (BioShop, prepared in DMSO), cefoxitin (Sigma, prepared in DMSO), cefsulodin (MP Biomedicals, 2 mg/mL), ceftazidime (Sigma, 4 mg/mL), cloxacillin (Sigma, prepared in DMSO), doripenem (Sigma), imipenem (AK Scientific, prepared in DMSO), methicillin (Cayman Chemicals), oxacillin (Cayman Chemicals, prepared in DMSO), penicillin G (Fluka Analytica), piperacillin (Sigma, 40 mg/mL), moenomycin complex (Caymen Chemicals, 0.01 mg/mL, prepared in DMSO), and vancomycin (Sigma, prepared in DMSO) were prepared at 10 mg/mL in sterile de-ionized H_2_O, except where noted, and stored at -20°C. Targocil was a gift from the Brown Lab and prepared at 5 mg/mL in DMSO. Lysostaphin from *Staphylococcus simulans* was purchased from Sigma Aldrich and prepared at 10 mg/mL in sterile de-ionized H_2_O.

### Inhibitory concentration assays

All bacterial strains were inoculated from -80°C glycerol stocks into 5 mL MHB and incubated with shaking, 200 rpm, 37°C for 16 h. Cultures were sub-cultured in a 1:500 dilution, grown for 4 h, and diluted in fresh MHB to an OD_600_ of ∼ 0.1, then diluted 1:500. Minimal inhibitory concentration values were determined as previously described, using compounds serially diluted in DMSO or sterile water ^27^. All MIC values are based on three independent experiments preformed in triplicate.

### Checkerboard assays

Checkerboard assays were set up and fractional inhibitory concentration indices (FICIs) were calculated as previously described ^27^. Briefly, 2 µL of compounds were added to an 8 x 8 section of a 96-well plate with increasing concentrations of one compound across the X-axis and increasing concentrations of the other compound across the Y-axis. Plates were incubated with shaking, 200 rpm, 37°C, 16 h and OD_600_ was measured and used to determine the MIC. Three biological replicates were averaged for each checkerboard.

### Transmission electron microscopy

*S. aureus* USA300 and USA300 Δ*tarO* were streaked on Mueller-Hinton agar (MHA) from -80°C glycerol stocks, then incubated at 37°C for 16 h. Single colonies were inoculated into 5 mL MHB and incubated with shaking, 200 rpm, 37°C for 16 h, then diluted 1:50 in fresh MHB and incubated for an additional 1.5 h. Cultures were normalized to OD_600_ ∼ 0.3 and incubated with DMSO, 8X MIC targocil, or 8-16X MIC BAY 11-7082 for 3 h. Cells were pelleted, washed with 100 µL MHB, then washed with 100 µL 2% glutaraldehyde fixative in phosphate buffer before being resuspended in 250 µL fixative and submitted to the Canadian Centre for Electron Microscopy for processing and sectioning. Cells were imaged using a JEOL 1200EX microscope. Images were taken at 75,000 x magnification (**Figure 6**) or 100,000 x magnification (**Figure S5**).

### Fluorescence Microscopy

To study PBP2 localization by fluorescence microscopy, COL sfGFP-PBP2 cultures were grown to mid-exponential phase (OD_600nm_ of 0.5 to 0.7) and 1 mL was collected and further incubated in the absence or presence of BAY 11-7082 (156 μM, 5X MIC) for 60 min at 37°C, before being collected by centrifugation. Cells were washed once with PBS, resuspended in 30 μL of PBS and 1 μL was placed on a thin layer of agarose (1.2 % in PBS). To evaluate septal enrichment of PBP2, the fluorescence ratio (FR) was calculated as the ratio of the median fluorescence of the 25% brightest pixels of the septum versus median fluorescence at the cell periphery, both corrected for background fluorescence. Microscopy was performed using a Zeiss Axio Observer microscope with a Plan-Apochromat 100X/1.4 oil Ph3 objective. Images were acquired with a Retiga R1 CCD camera (QImaging) a white-light source HXP 120 V (Zeiss) using Zen Blue software (Zeiss).

To determine the effect of antibiotics on *vraTSR* promoter activity strain JE2 Pvra-mNG was incubated in the presence or absence of BAY-11-7082 (156 mM, 5X MIC) and in the presence or absence of vancomycin (12.5 mg/ml, 10X MIC) for 60 min at 37°C. Pairs of samples were then imaged on the same microscopy slide. For this, JE2 Pvra-mNG cells not exposed to a antimicrobial compound were labeled with DNA dye Hoechst 33342 (1 μg mL^−1^) while cells exposed to the antimicrobial compound were not labelled. Both samples were washed with PBS, and then the two samples were mixed prior to visualization by epifluorescence microscopy as described above. Total fluorescence of each cell was calculated using eHooke software version 1.1, as previously described ^51^.

## Supporting information

Coles et al Supplementary Figures S1-S5

## ACKNOWLEDGEMENTS

We thank the Brown lab for providing targocil and the Δ*tarO* strain of MRSA, Andreia Duarte for the construction of the pPvra-mNG plasmid and Eric Brown, Shawn French, and Maya Farha for helpful ideas and discussions. We thank Cezar Khursigara and Erin Anderson for helpful discussions about electron microscopy and Marcia Reid from the Canadian Centre for Electron Microscopy for sample preparation and microscopy imaging. This work was supported by a Discovery Grant to LLB from the Natural Sciences and Engineering Research Council of Canada [RGPIN- 2021-04237] and by the European Research Council through ERC Advanced Grant [101096393] to MGP. LLB holds a Tier 1 Canada Research Chair in Microbe-Surface Interactions [CRC-2021-00103]. VEC held an Ontario Graduate Scholarship.

