## Supplementary material for "BAY 11-7082 potentiates select β-lactams to inhibit growth of methicillin-resistant *Staphylococcus aureus*": Coles et al Supplementary Figures S1-S5

**SUPPORTING INFORMATION for: BAY 11-7082 potentiates select β-lactams to inhibit growth of methicillin-resistant *Staphylococcus aureus*.'** Victoria E. Coles[1], Patricia Reed[2], Mariana G. Pinho[2], and Lori L. Burrows[1]. [1] Department of Biochemistry and Biomedical Sciences and the Michael G. DeGroote Institute for Infectious Disease Research, McMaster University, Hamilton, ON and [2] Instituto de Tecnologia Química e Biológica António Xavier, Universidade NOVA de Lisboa, Oeiras, Portugal.

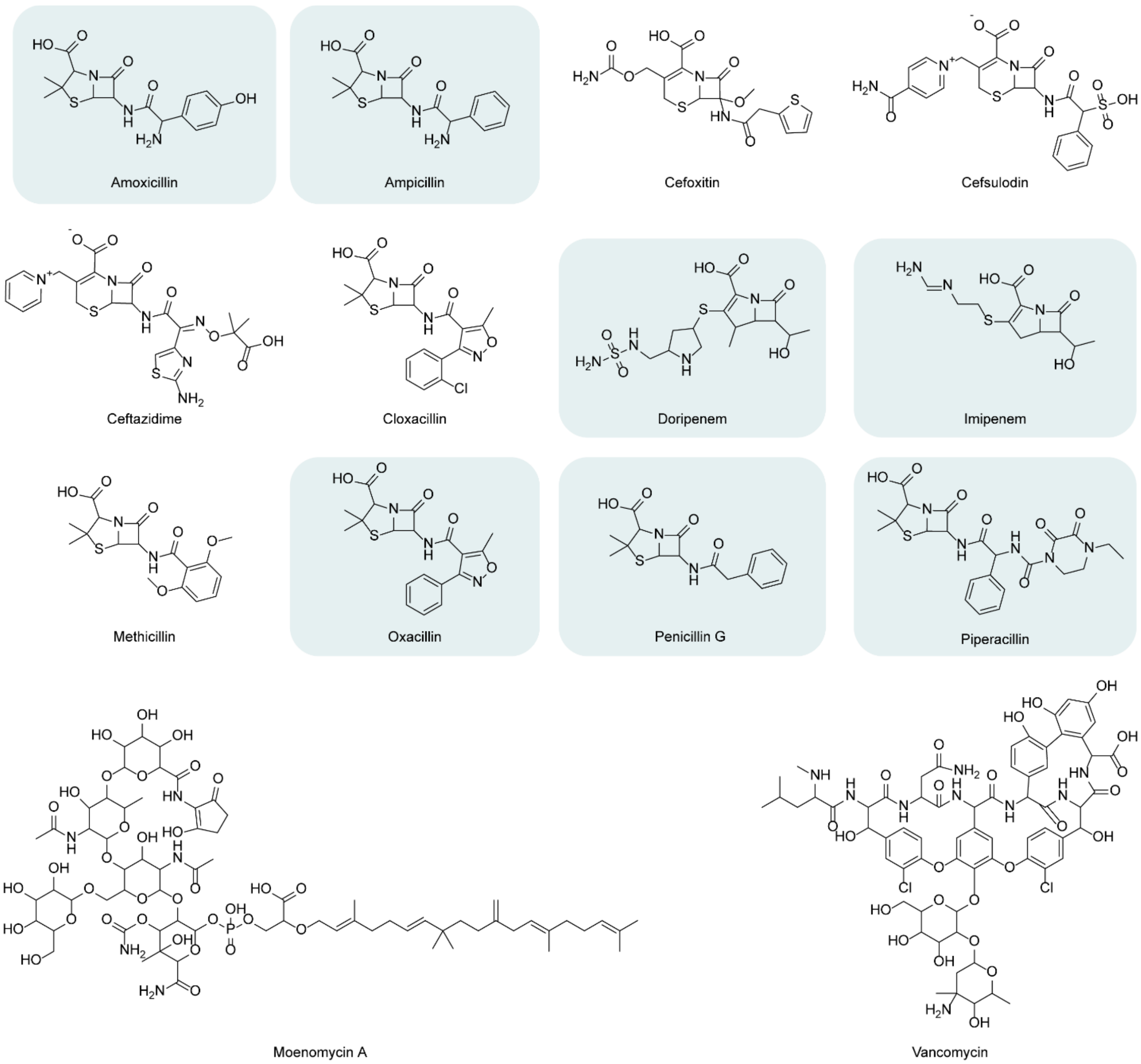


**Figure S1. Structures of β-lactams and non-β-lactam peptidoglycan-targeting antibiotics tested for activity against *S. aureus* USA300 (MRSA) and ATCC 29213 (MSSA) in combination with BAY 11-7082.** Compounds that synergize with BAY 11-7082 are highlighted.

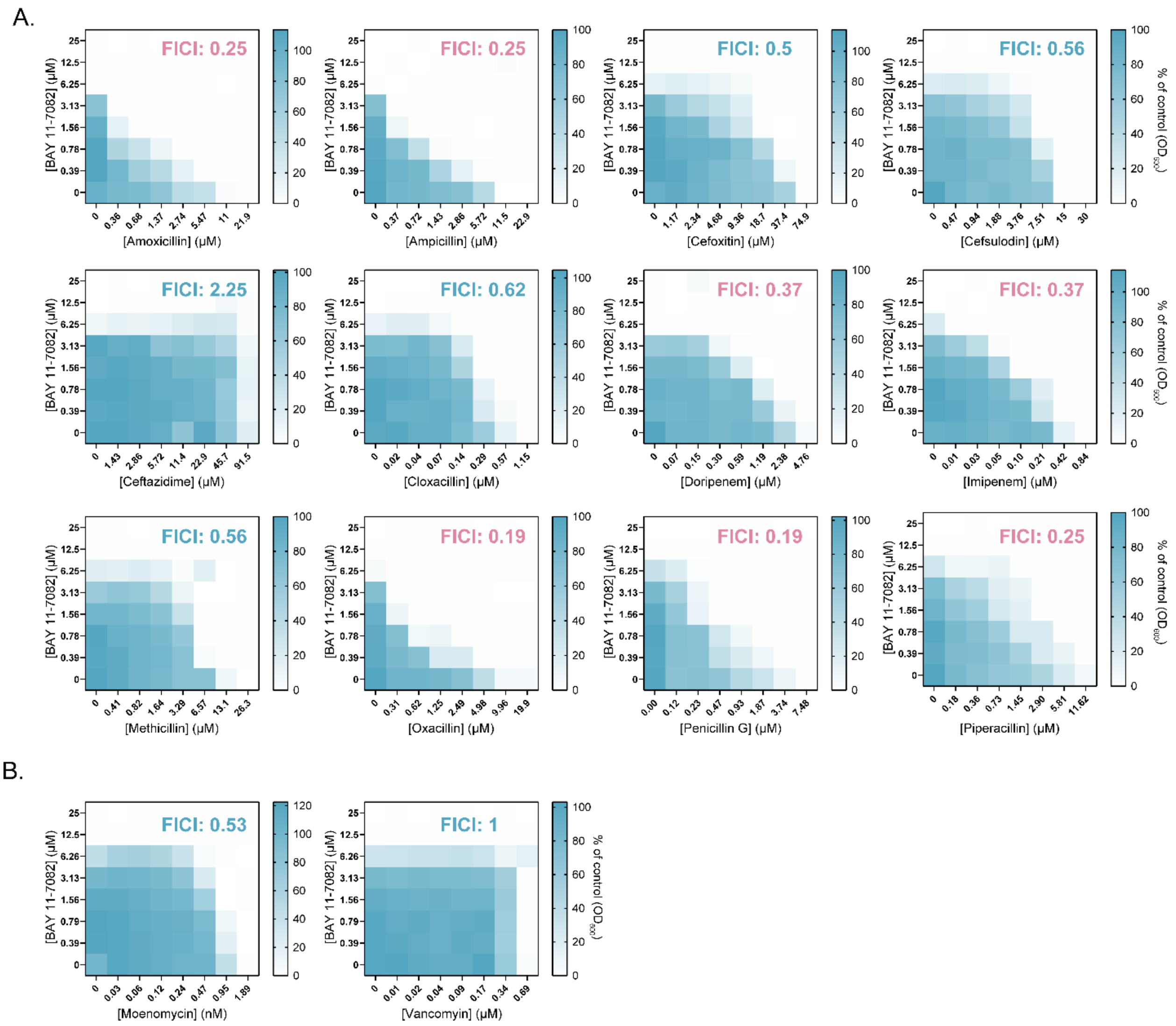


**Figure S2. Checkerboards of *S. aureus* USA300 (MRSA) grown with increasing concentration of BAY 11-7082 and A. a β-lactam or B. other cell-wall active antibiotic, where synergy (FICI < 0.5) is shown in pink text while indifference (FICI ≥ 0.5) is shown in blue.** Data represent an average of three independent experiments.

A.

B.

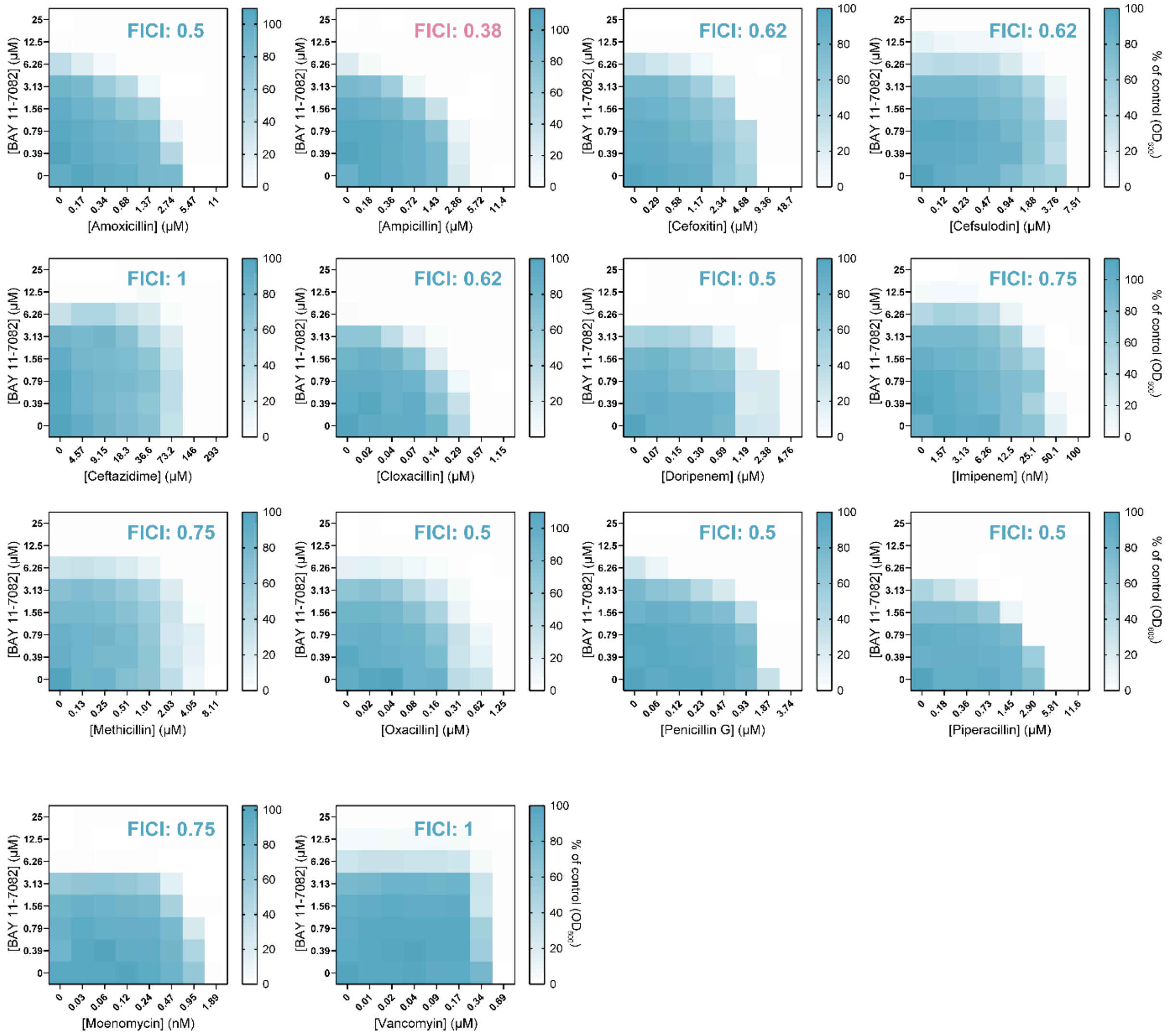


**Figure S3. Checkerboards of *S. aureus* ATCC 29213 (MSSA) grown with increasing concentration of BAY 11-7082 and A. a β-lactam or B. other cell-wall active antibiotic, where synergy (FICI < 0.5) is shown in pink text while indifference (FICI ≥ 0.5) is shown in blue.** Data represent an average of three independent experiments.

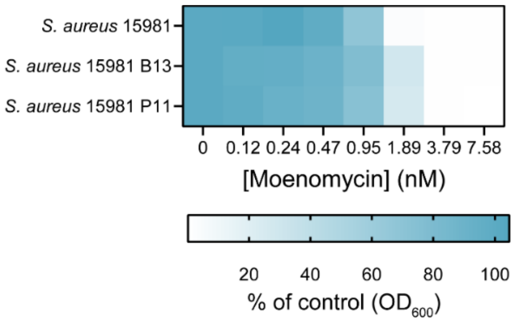


**Figure S4. Planktonic growth (optical density at 600 nm as a percent of the vehicle control) of *S. aureus* 15981 (MRSA), the BAY 11-7082-resistant mutant B13, and the PSPC-resistant mutant P11.** Cells were grown in the presence of increasing concentrations of the antibiotic moenomycin (nM). Data represent an average of three independent experiments performed in triplicate.

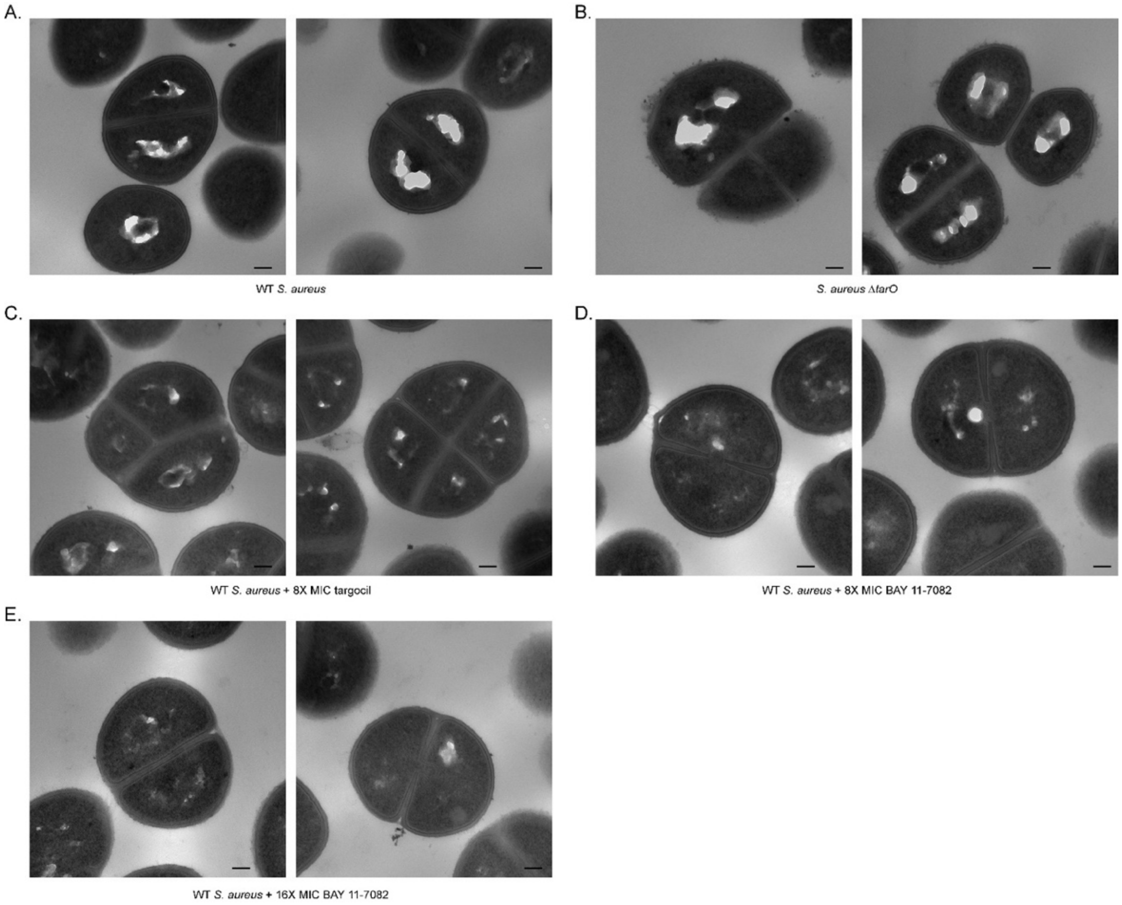


**Figure S5. Additional fields of view of electron micrographs. A.** Untreated wild type *S. aureus* USA300. **B.** *S. aureus* USA300 *ΔtarO* cells. **C.** Wild-type *S. aureus* USA300 treated with 33.6 µM targocil**. D.** Wild-type *S. aureus* USA300 treated with 50 µM BAY 11-7082**. E.** Wild-type *S. aureus* USA300 treated with 100 µM BAY 11-7082. Scale bar = 100 nm.
